# Cortical Surface-Informed Volumetric Spatial Smoothing of fMRI Data via Graph Signal Processing

**DOI:** 10.1101/2021.05.04.442605

**Authors:** Hamid Behjat, Carl-Fredrik Westin, Iman Aganj

## Abstract

Conventionally, as a preprocessing step, functional MRI (fMRI) data are spatially smoothed before further analysis, be it for activation mapping on task-based fMRI or functional connectivity analysis on resting-state fMRI data. When images are smoothed volumetrically, however, isotropic Gaussian kernels are generally used, which do not adapt to the underlying brain structure. Alternatively, cortical surface smoothing procedures provide the benefit of adapting the smoothing process to the underlying morphology, but require projecting volumetric data on to the surface. In this paper, leveraging principles from graph signal processing, we propose a *volumetric* spatial smoothing method that takes advantage of the gray-white and pial cortical surfaces, and as such, adapts the filtering process to the underlying morphological details at each point in the cortex.

## I. INTRODUCTION

Functional magnetic resonance imaging (fMRI) is a key non-invasive imaging modality for studying brain activity based on the blood-oxygen-level-dependent signal [1]. When analyzing fMRI data, a conventional preprocessing step is to spatially smooth the data, which is generally done for two reasons: (i) to deal with the multiple testings through resorting to Gaussian random field theory [2]—as opposed to using stricter methods such as the Bonferroni correction— and (ii) to improve the data signal-to-noise ratio (SNR) by employing spatial smoothing such as lowpass filtering. With regards to (i), alternative methods have been proposed that obviate the need for Gaussian smoothing, namely, but not limited to, false discovery rate correction [3], permutation testing [4], and wavelet denoising [5]. With regards to (ii), the conventional use of isotropic filters—such as the Gaussian filter—comes at the expense of a loss in fine spatial details of the underlying activity [6], since in virtue of the matched filter argument, spatial filters are optimal in denoising data only if their shape and size conform to the underlying signal of interest. Given that the spatial profile of brain fMRI data is confined by the underlying neuroanatomical structure, alternative spatial filtering methods that incorporate knowledge about the underlying brain structure have been proposed, broadly classified into surface-based—see e.g. [7], [8], [9], [10], [11]—and volume-based—see e.g. [12], [13], [14], [15]—methods, which differ from methods such as [16], [17], [18], [19] in that they leverage an independent contrast image—different from the data to be smoothed— that confines the spatial profile of the filters.

In this paper, we propose a scheme that enables volumetric spatial smoothing of cerebral cortical fMRI data in such way that filtering is informed by the morphological structure of the cerebral cortex provided by cortical surface extractions. At the heart of the method lies the representation of a cerebral hemisphere cortex—a.k.a. the cortical ribbon—as a graph [20], [21], wherein each voxel is represented as a vertex and edges are defined based on the Euclidean adjacency of voxels but constrained by topological information provided by reconstructed white-gray border and pial cortical surfaces [22]. The graphs are subject-specific and thus encode the unique morphology of each individual’s cortical structure. Principles from the recently emerged field of graph signal processing [23], [24] are then leveraged to perform spatial filtering of fMRI data in such way that the filter profiles are adapted to cortical morphology.

## II. METHODS

### A. Graph signal processing: fundamentals

An undirected, non-weighted graph, denoted as 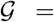 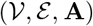, consists of a set 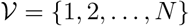 of *N* vertices and a set 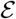 of edges (i.e., pairs (*i, j*) where 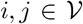), and can be represented by an *N × N* symmetric matrix **A**, i.e., the graph adjacency matrix, with its elements defined as *a*_*i,j*_ = 1 if 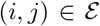 and *a*_*i,j*_ = 0 otherwise. The graph’s symmetric normalized Laplacian matrix, denoted as **L**, is given as 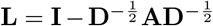, where **D** denotes the graph’s degree matrix, which is a diagonal matrix with elements given as *d*_*i,i*_ = Σ_*j*_*a*_*i,j*_, and **I** denotes the identity matrix. Since **L** is real, symmetric, diagonally dominant, and with non-negative diagonal entries, it is positive semi-definite, and therefore, it can be eigen-decomposed as

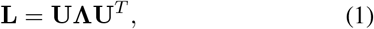

where **Λ** is a diagonal matrix that stores the eigenvalues *λ*_1_*, …, λ*_*N*_ ≔ *λ*_max_ and **U** is a matrix that stores the corresponding eigenvectors in its columns, **U** = [**u**_1_|**u**_2_| · · · |**u**_*N*_]; hereon, we refer to eigenvectors of **U** as eigenmodes, in line with the convention in the neuroimaging community. The eigenvalues of **U** define a spectrum for 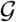 [25], a space that can be seen as an extension of the Euclidean Fourier domain.

Given a graph 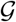, data residing on the vertices of 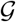 can be seen as a *graph signal*, denoted as **f** ∈ ℝ^*N*^. Using **U**, **f** can be transformed to the spectrum 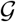, denoted as 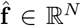 and obtained as 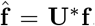, which is a representation that is commonly referred to as the *graph Fourier transform* of **f**. This transformation satisfies Parseval’s energy conservation relation [26], i.e., 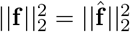.

Given a graph 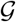 and a graph signal **f**, it is beneficial to be able to implement filtering operations on **f** as in conventional signal processing. In particular, it is of interest to define graph filters that entail specific spectral properties, for instance, defining lowpass and highpass filters as in conventional signal processing. A graph filter can be conveniently defined in the graph Laplacian spectral domain as *h* : [0, *λ*_max_] → ℝ, using which, filtering of **f**, denoted as **f**_*h*_, can be straightforwardly implemented in the spectral domain by modulating 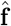 with *h*(·), and then transforming the modulated signal to the vertex domain, given as

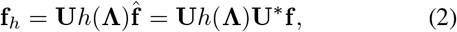

where *h*(**Λ**) denotes a diagonal matrix with its entries computed by sampling *h*(·) at the eigenvalues. However, a shortcoming of this approach to graph signal filtering is that it requires **Λ** and **U**, i.e., eigen-decomposition of **L**. This is computationally highly impractical for large graphs, and, in particular, infeasable for the graphs proposed in this work that are typically of sizes in the range 100 K to 200 K vertices. An alternative approach is to approximate *h*(·) as a polynomial, denoted as 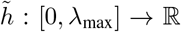, and to then implement filtering within the vertex domain as [27]

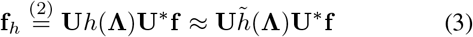

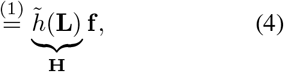

where in the last equality we used the property 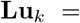 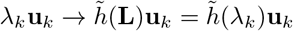. As such, graph signal filtering can be seamlessly implemented through the mere use of polynomial matrix operations on **L**. In particular, it is insightful to observe that each row *k* in 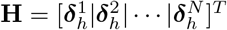, where 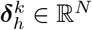, is the vertex representation of spectral kernel *h*(·) when instantiated at vertex *k*; these set of vectors provide the impulse response set for spectral kernel *h*(·), which, unlike that in conventional signal processing, are *shift-variant*.

### B. Dataset

We used the Human Connectome Project (HCP) dataset [28] in this study; the acquisition of the HCP dataset was approved by the Washington University Institutional Review Board and informed consent was obtained from all subjects. In particular, we used a subset of the dataset, the 100 unrelated subjects (54% female, mean age = 29.11± 3.67, age range = 22-36). To construct cortical graphs, we used the minimally preprocessed structural data, which come at 0.7 mm isotropic resolution, and the associated extended preprocessed version of the data, which included FreeSurfer surface extraction. We also used the HCP functional task fMRI data, which come at 2 mm isotopic resolution, and consisted of seven functional tasks: Emotion, Gambling, Language, Motor, Relational, Social, and Working Memory. The spatial smoothing methodology proposed in this work heavily relies on accurate co-registration between the structural and functional data, which is already implemented in the HCP preprocessed data. A full description of the imaging parameters and prepocessing steps can be found in [29].

### C. Cerebral hemisphere cortex (CHC) graphs

We design subject-specific, voxel-resolution graphs that encode the morphology of CHC using the method presented in [20], [21], which is based on leveraging the volumetric representation of a subject’s CHC as extracted by FreeSurfer^1^ [22] from a T1-weighted MRI image, the so called *ribbon* representation. Each vertex of the 3D ribbon is considered as a graph vertex. Graph edges are preliminarily defined based on the adjacency of voxels within the cubic lattice of size 3×3×3, i.e., 26 neighborhood connectivity. Consequently, spurious edges that are anatomically unjustifiable—e.g. edges that connect voxels that lie on opposite sides of narrow sulci—are pruned out by using pial and white surface representations extracted using FreeSurfer. For a more detailed description of the design, we refer the interested reader to [20], [21].

### D. Spectral graph heat kernel filters

Given a CHC graph, we design spatial smoothing filters associated with a heat kernel profile defined on the graph spectrum, i.e., *k*(*λ*) = *e*^−*τλ*^,*⩝λ* ∈ [0, *λ*_max_], where *τ* is a free parameter that specifies the resulting filter’s spatial extent; see Fig. 1. For a given *τ*, we approximated *k*(*λ*) by a Chebyshev polynomial—approximating a minimax polynomial, minimizing an upper bound on the approximation error [27]—and applied filtering as in (4).

**Fig. 1.**
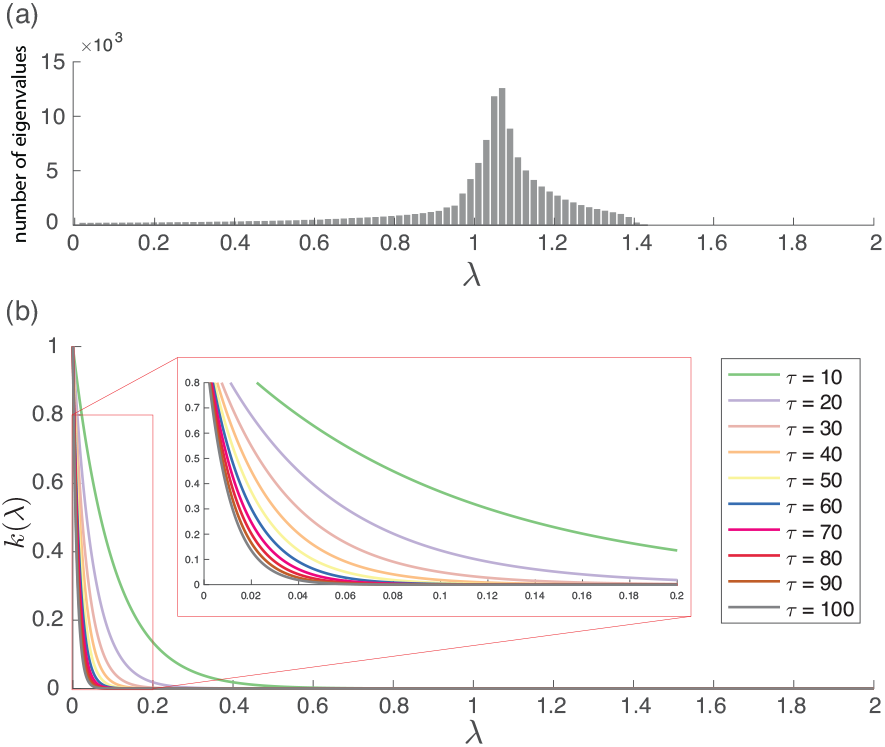
(a) Distribution of graph Laplacian eigenvalues of a representative subject’s left hemisphere CHC graph. (b) Spectral graph heat kernels.

### E. Construction of cortical activity phantoms

A semi-synthetic phantom dataset was constructed through synthesizing cortical activation patterns and corrupting them with white Gaussian additive noise. The anatomical scans of the first 10 subjects from the HCP dataset were used to generate activation patterns that have confounding spatial shapes that are constrained by the cortical morphology of each individual subject, using a method similar to that proposed in [12]. Specifically, for each subject *i*, 10 random seed points were selected from within the cortical ribbon, treated as distinct centers of functional activity. The points were then represented as an indicator vector **x**_*i*_ ∈ ℝ^*N*^, wherein **x**_*i*_[*n*] = 1 if *n* corresponds to a center point and **x**_*i*_[*n*] = 0 otherwise, i.e., ||**x**_*i*_||_1_ = 10. An activation pattern that diffuses from the center points along the subject’s cortex was generated as **y**_*i*_ = **z**_*i*_/ max_*n*_{**z**_*i*_[*n*]}, where 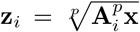, and **A**_*i*_ denotes the adjacency matrix of subject *i*, and *p* is parameter that determines the extent of the diffused pattern. In all the following analyses, *p* was set to 5. The resulting activation patterns are unique to each subject and exhibit non-binary distributions of value within the range [0, 1]. The vector-formed activity patterns 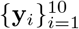 were then placed back into 3D volumetric representations; Fig. 2 shows the resulting phantoms across the 10 subjects. The phantoms were then corrupted with additive white Gaussian noise; phantoms with four noise levels were generated—standard deviations *σ*_*n*_ = 2, 4, 8 and 16, resulting in contrast-to-noise ratios (CNR)—max_*n*_{**y**_*i*_}/*σ*_*n*_ [30]—of 1/2, 1/4, 1/8 and 1/16, respectively. Ten realizations were generated for each noise level, resulting in a total of 100 signals for each CNR.

**Fig. 2.**
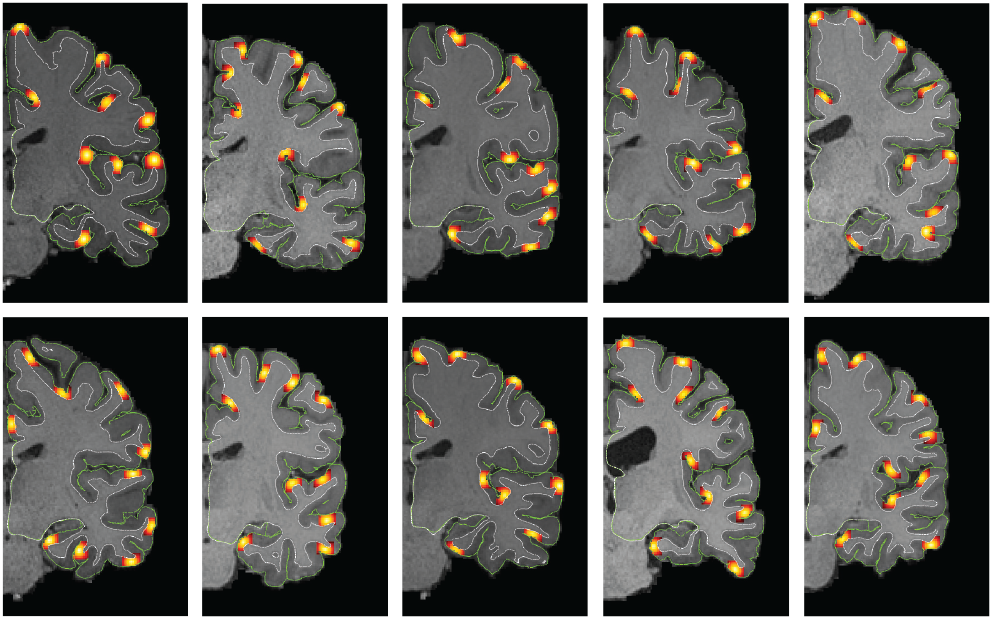
Simulated cortical activity phantoms on 10 HCP subjects.

## III. RESULTS AND DISCUSSION

Fig. 3 shows realizations of a representative set of graph spatial smoothing filters when localized at different parts along the cortical ribbon. In contrast to conventional Gaussian smoothing wherein the spatial shape of the filtering kernel is the same at each position across the image, in graphbased spatial smoothing, the shape of graph filters adapts to each specific position along the domain they are realized at, based on the encoding provided by the graph.

**Fig. 3.**
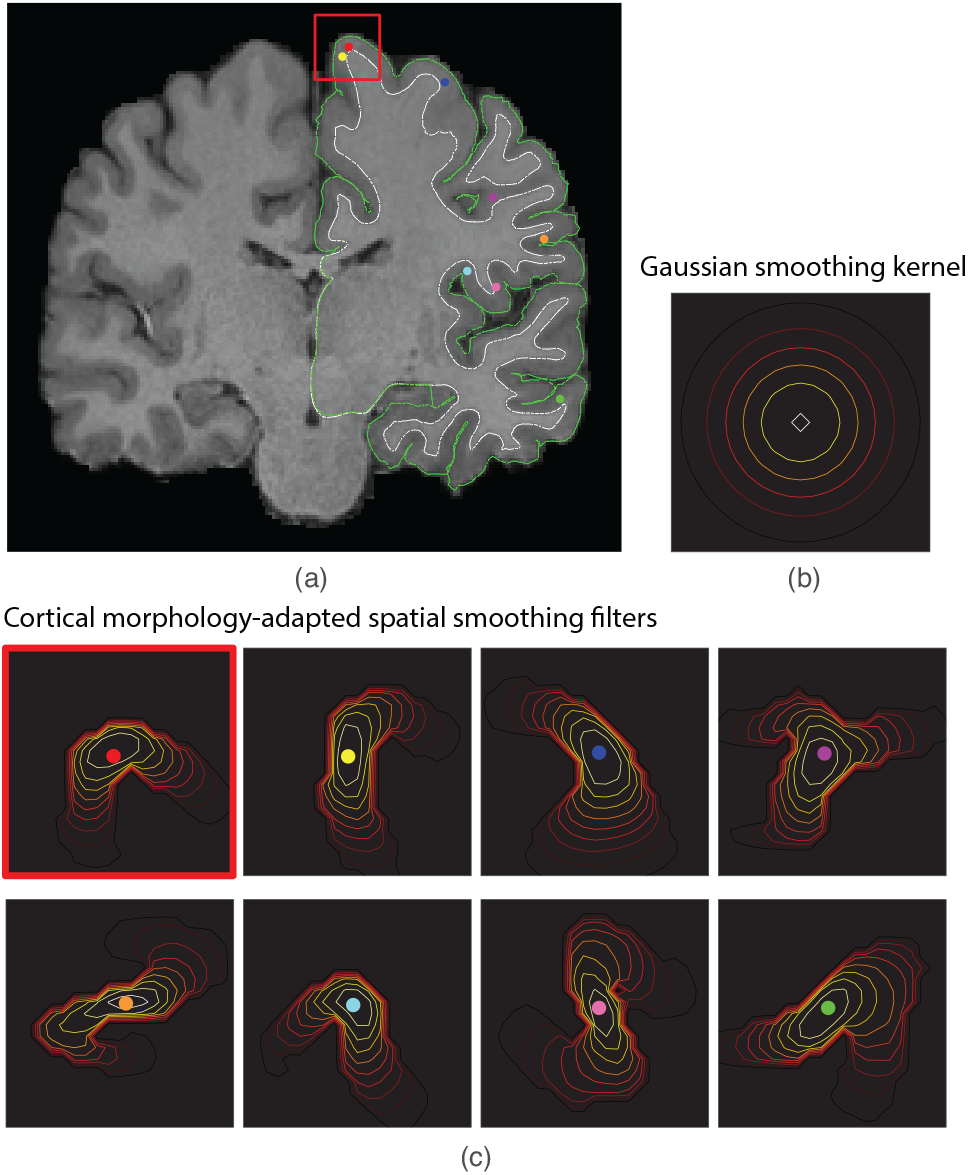
(a) Voxels that fall within the cerebral cortex—as defined by the region in between the pial (green) and white (white) surfaces—of a given hemisphere define the vertices of a CHC graph, and the graph edges are defined based on geodesic adjacency of voxels within their 3 × 3 × 3 voxel neighborhood [20], [21]. (b) When performing volumetric smoothing of fMRI data, Gaussian kernels are conventionally used, which are isotropic and not adapted to the underlying cortical morphology; contours of a Gaussian kernel with FWHM = 8 mm is displayed. (c) Using CHC graphs, spatial filters that adapt to the underlying cortical morphology are designed; in particular, a unique spatial filter is obtained at each position (voxel) within the cerebral cortex. Contours of eight representative graph filters associated to a heat kernel spectral profile, with parameter *τ* = 40, are displayed.

To validate the strength of the proposed filters in denoising fMRI data, receiver operating characteristic (ROC) analysis was performed on the phantom set, see Section II-E. Each noise-added phantom was spatially smoothed with <u>Gau</u>ssian <u>s</u>patial <u>s</u>moothing (GAUSS) and <u>gra</u>ph-based <u>s</u>patial <u>s</u>moothing (GRASS) over a range of filter sizes, FWHM = 1, 2, … 14 and *τ* = 10, 15, …, 100, respectively. For GAUSS, two approaches were compared: unconstrained GAUSS (uGAUSS) and constrained GAUSS (cGAUSS). In uGAUSS, the data were smoothed without masking out regions outside the ribbon; that is, the filter has access to values not only within the cortical ribbon but also to those in adjacent white matter or CSF. In cGAUSS, the region outside the ribbon was masked out (set to zero), and consequently, normalized convolution was performed; that is, filtering was done in three steps, first, the masked data was smoothed, then, a mask representing the cortical ribbon (with values one inside the ribbon and zero outside) was smoothed, and consequently, the former was divided by the latter.

After filtering, smoothed volumes were thresholded at 100 uniformly-spaced consecutive levels spanning the minimum and maximum value in each filtered volume to generate ROC curves—wherein the clean phantoms were treated as ground truth, and in turn, the area under curve of each ROC curve (AUCROC) was computed. Fig. 4 shows the results. Between the two GAUSS approaches, cGAUSS outperformed uGAUSS as it uses additional information about the delineation of the cortical ribbon, preventing mixture of pure noise from surrounding white matter and CSF. Across CNRs, GRASS showed superior performance over GAUSS. Although both cGAUSS and GRASS exploit cortical surface information, GRASS outperforms cGAUSS since i) it leverages filters whose spatial profile adapts to the domain, whereas in cGAUSS, the filters are merely masked by the cortical ribbon, and ii) it prevents mixing of values at touching parts of the cortex that are only adjacent in Euclidean sense but not in geodesic sense, for instance touching banks of narrow sulci. Best performance for GAUSS was obtained for FWHMs within the range 6 to 10, and for GRASS it was obtained for *τ* within the range 30 to 50.

**Fig. 4.**
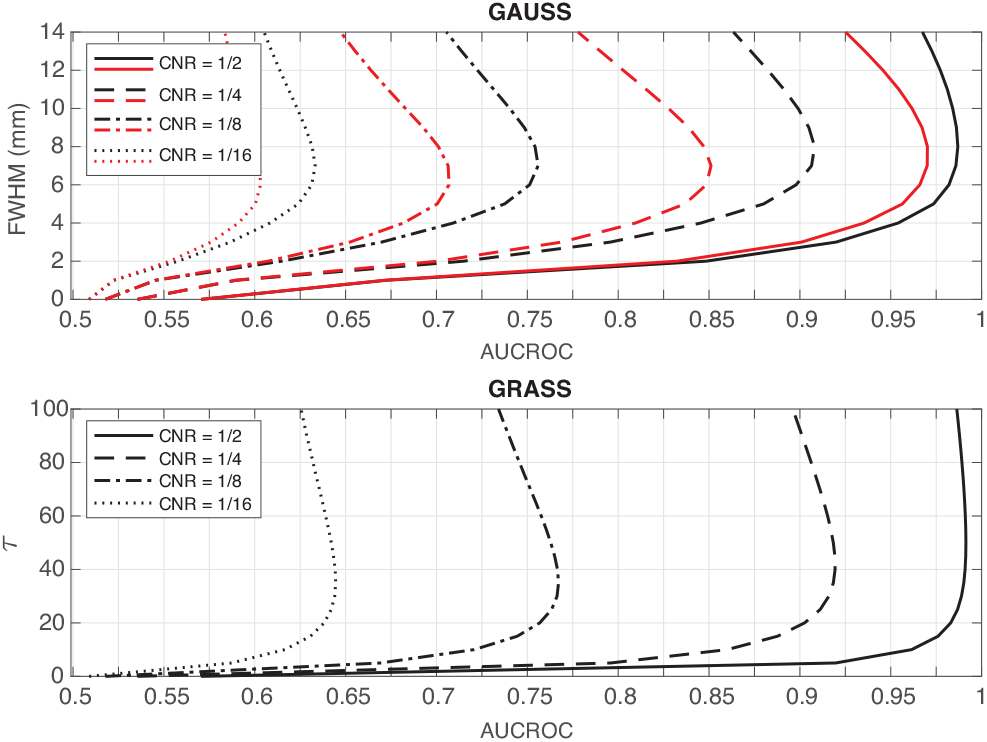
Each point along the curves is the average AUCROC over 100 realizations (10 phantoms × 10 realizations of noise). For GAUSS, the curves shown in red and black are for uGAUSS and cGAUSS, respectively.

We then performed Dice analysis to quantify the extent of difference between detection maps resulting from GRASS and cGAUSS on the real data. Given the large number of available combinations of cGAUSS and GRASS for different filter sizes, and lack of a one-to-one mapping between a Gaussian kernel of a given FWHM and a graph filter of a specific *τ*, we selected a subset of filter sizes for cGAUSS and GRASS for the analysis performed on the real data, based on the AUCROC performances obtained on the simulated data. In particular, we selected five filter sizes that provided the overall best performance for cGAUSS (FWHM = 6, 7, 8, 9 and 10) and GRASS (*τ* = 30, 35, 40, 45 and 50); see Fig. 4. With this selection of filter sizes, we are implicitly assuming that underlying activations in the real data have spatial spreads that are of approximately the same size as those in the simulated data. For each subject, and each functional task, the fMRI data were smoothed using the 10 above-listed filter settings, and activation mapping was then performed for one of the experimental conditions of each task (Emotion: *fear*, Gambling: *win*, Language: *math*, Motor: *left hand*, Relational: *match*, Social: *mental*, Working Memory: *body 0-back*; for details about the tasks and each experimental condition, we refer the interested reader to [31]). Resulting t-maps were then thresholded at 5% false discovery rate. For a given subject, a given task, and a given *τ*, the Dice similarity between the associated GRASS detection map and the five detection maps for cGAUSS were computed as *d*_*τ,fwhm*_ = 2|*M*_*τ*_⋂*M*_fwhm_|/|*M*_*τ*_ + |*M*_fwhm_, for FWHM = 6, …, 10, where *M*_fwhm_ and *M*_*τ*_ denote the set of voxels in the detected map for cGAUSS and GRASS, respectively, and | · | denotes set cardinality. The maximum Dice was then selected, i.e., max{*d*_*τ*,fwhm_}_fwhm=6,···,10_. This analysis was repeated across subjects, tasks, and the five *τ* values. If activation mapping on data smoothed with neither cGAUSS nor GRASS resulted in any detections, Dice similarity was not computed.

Fig. 5(a)-left shows the maximum Dice for different tasks for *τ* = 40, showing values within the range 0 to 0.92 and an average of 0.8 across the seven tasks^2^. A similar average Dice similarity was observed using the other four *τ* values; see Fig. 5(a)-right. Despite the similarity of detection maps for GRASS and cGAUSS, the two approaches nevertheless resulted in a notable set of detections that are not common between the two. Figs. 5(b) and (c) show the extent of unique detections by cGAUSS and GRASS, respectively, as ratios of total number of detections by each method; in particular, Fig. 5(b) shows |*M*_fwhm_| − |*M*_τ_ ⋂ *M*_fwhm_|, and Fig. 5(c) shows |*M*_τ_| − |*M*_τ_ ⋂ *M*_fwhm_|, for the *τ* and the FWHM for which Dice values are shown in Fig. 5(a). On average, 21%^3^ of the detections by cGAUSS were not present in the detection maps for GRASS, and 17%^4^ vice versa. Due to lack of ground truth, it is not possible to make any concrete statements when interpreting these results. However, if we are to accept the validity of the simulation results— specifically, that the generated synthetic phantoms resemble true activations—the superior performance of GRASS over cGAUSS on the simulations suggests the potential of having obtained a better specificity and sensitivity using GRASS.

**Fig. 5.**
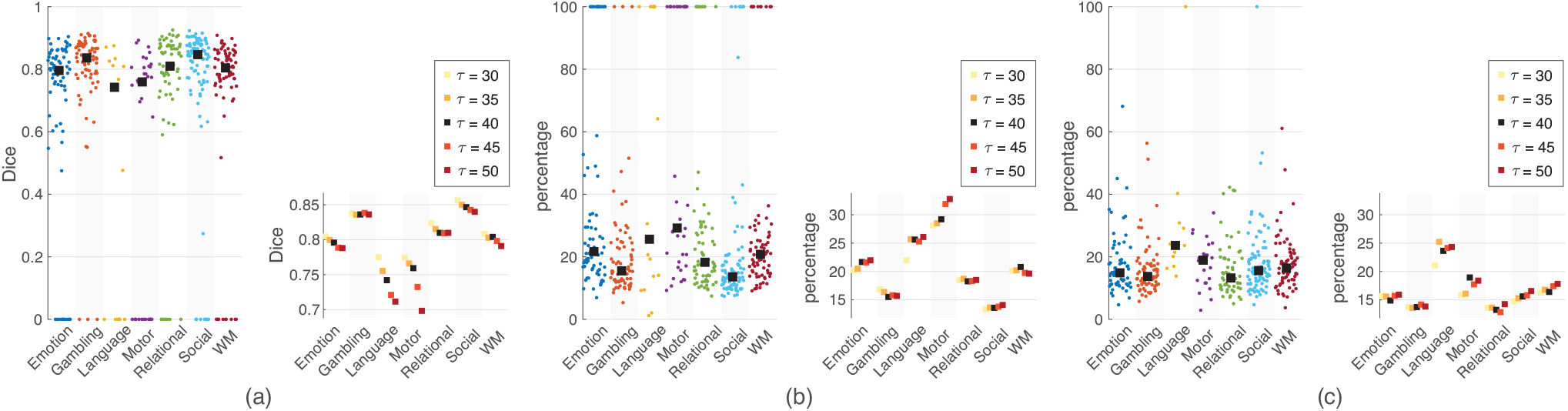
(a) Left: maximum Dice similarity between detection map resulting from data smoothed with GRASS with *τ* = 40 and 5 detection maps resulting from data smoothed with cGAUSS, FWHMs = 6, 7, 8, 9 and 10. Each dot represents one subject. The mean value across subjects is shown by ■. Right: mean values same as in the left plot but for other *τ* values. (b) Left: for the pair of *τ* and FWHM for which Dice values are shown in (a), percentage of detections for cGAUSS that are not present in the detection map for GRASS, i.e., unique detections for cGAUSS. Each dot represents one subject. The mean value across subjects is shown by ■. Right: mean values similar to the left plot but for other *τ* values. (c) Same as in (b) but for GRASS.

## IV. CONCLUSIONS

We proposed a method that enables volumetric spatial smoothing of functional data in such way that filtering is informed by the morphological structure of the cerebral cortex, as defined by extracted cortical surfaces. In future work, we will compare the proposed method with spatial smoothing performed on cortical surface [10], [14], and will also explore the benefits of the proposed method for processing high spatial resolution fMRI data [32], [33].

## Footnotes

1 https://surfer.nmr.mgh.harvard.edu

2 Emotion: 0.8, Gambling: 0.84, Language: 0.74, Motor: 0.76, Relational: 0.81, Social: 0.85 and WM: 0.8.

3 Emotion: 22%, Gambling: 15%, Language: 26%, Motor: 29%, Relational:18%, Social:14% and WM: 21%.

4 Emotion: 15%, Gambling: 14%, Language: 24%, Motor: 19%, Relational: 13%, Social: 16% and WM: 16%.

